## Supplemental Tables for "Dysfunctional Mitochondria in Cardiac Fibers of a Williams-Beuren Syndrome Mouse Model"

### Supplemental Table S1: Statistical data of Oxygen consumption and ATP production

| **Unpaired t-test- Multiple comparisons Holm-Sidak method** | | | | |
|---|---|---|---|---|
| | **t-ratio** | **df** | **P value** | **Adjusted P value** |
| LEAK | 1.88927 | 10 | 0.088168 | 0.088168 |
| OXPHOS CI | 2.66538 | 10 | 0.023680 | 0.046799 |
| OXPHOS CI+CII | 3.63116 | 10 | 0.004604 | 0.013748 |
| OXPHOS CIV | 8.10377 | 9 | 0.000020 | 0.000100 |
| ETC CII | 4.37771 | 10 | 0.001382 | 0.005518 |
| **Unpaired t-test** | | | | |
| | **t-ratio** | **df** | **P value** | |
| RCR | 3.470 | 10 | 0.0060 | |
| MEFs-ATP | 4.315 | 4 | 0.0125 | |

### Supplemental Table S2: Statistical data of genomic copy number and OXPHOS complex

| **Unpaired t-test: genomic copy number** | | | | |
|---|---|---|---|---|
| | **t-ratio** | **df** | **P value** | |
| Cardíac fibers | 2.658 | 9 | 0.0261 | |
| MEFs | 4.066 | 8 | 0.0036 | |
| **Unpaired t-test- Multiple comparisons Holm-Sidak method** | | | | |
| | **t-ratio** | **df** | **P value** | **Adjusted P value** |
| CI-NDUFB8 | 5.038 | 7 | 0.0015 | 0.0075 |
| CII-SDHB | 4.021 | 7 | 0.0050 | 0.0125 |
| CIII-UQCRC2 | 4.425 | 7 | 0.0031 | 0.0122 |
| CIV-MTCO1 | 3.051 | 6 | 0.0225 | 0.0225 |
| CV-ATP5A | 4.169 | 7 | 0.0042 | 0.0125 |

**Supplemental Table S3: Statistical data of mitochondrial morphology**

| **Unpaired t-test** | | | |
|---|---|---|---|
| | **t-ratio** | **df** | **P value** |
| Density | 2.626 | 6 | 0.0393 |
| Average Size | 3.321 | 6 | 0.0160 |
| Circularity | 2.865 | 6 | 0.0286 |

**Supplemental Table S4: Statistical data of mitochondrial dynamics**

| **Unpaired t-test- Multiple comparisons Holm-Sidak method** | | | | |
|---|---|---|---|---|
| | **t-ratio** | **df** | **P value** | **Adjusted P value** |
| L-OPA1 | 3.387 | 12 | 0.005400 | 0.010632 |
| S-OPA1 | 3.394 | 12 | 0.005330 | 0.010632 |
| **Unpaired t-test** | | | | |
| | **t-ratio** | **df** | **P value** | |
| FIS1 | 4.887 | 7 | 0.0018 | |
| MTF1 | 3.859 | 8 | 0.0048 | |
| MTF2 | 0.205 | 7 | 0.8431 | |

**Supplemental Table S5: Primer sequences for genotyping and genomic copy number**

| **Gene** | **Sequence** | **Tm ºC** |
|---|---|---|
| **HPRT** | 5’-CTCTGAGGCTTCAAAGGTTC-3’ | 56.7 |
| | 5’-AATCCAGCTTGTTTGGGCTA-3’ | 59.7 |
| **TRDC** | 5’-CAAATGTTGCTTGTCTGGTG-3’ | 57.7 |
| | 5’-GTCAGTCGAGTGCACAGTTT-3’ | 56.4 |
| **GADPH** | 5′-ATGACTCCACTCACGGCAAAT-3′ | 61.9 |
| | 5′- GGGTCTCGCTCCTGGAAGAT-3′ | 63.0 |
| **mt-ND1** | 5’-GGATCCGAGCATCTTATCCA-3’ | 60.0 |
| | 5’- GGTGGTACTCCCGCTGTAAA-3’ | 60.0 |
